# Mathematical Modeling Reveals the Factors Involved in the Phenomena of Cancer Stem Cells Stabilization

**DOI:** 10.1101/475285

**Authors:** N. Bessonov, G. Pinna, A. Minarsky, A. Harel-Bellan, N. Morozova

**Affiliations:** Institute of Problems of Mechanical Engineering, Russian Academy of Sciences, St-Petersburg, Russia; Institute for Integrative Biology of the Cell (I2BC), CEA, CNRS, University Paris-Sud, University Paris-Saclay, 91198, Gif-sur-Yvette cedex, France; St-Petersburg Academic University, Russian Academy of Sciences, St-Petersburg, Russia; Institut des Hautes Etudes Scientiques, Bures-sur-Yvette, France

## Abstract

Cancer Stem Cells (CSC), a subset of cancer cells resembling normal stem cells with self-renewal and asymmetric division capabilities, are present at various but low proportions in many tumors and are thought to be responsible for tumor relapses following conventional cancer therapies. In vitro, most intriguingly, when isolated, CSCs return to their original proportion level as shown by various investigators. This phenomenon still remains to be explained.

We suggest a mathematical model of cancer cell population dynamics, based on the main parameters of cell population dynamics, including the proliferation rates, the rates of cell death and the frequency of symmetric and asymmetric cell divisions both in CSCs and in non-CSCs. This model should help elucidating some important factors underlying the dynamics of the two populations, first of all, the phenomena of cancer stem cell population stabilization.

**Author Summary:** Cancer Stem Cells (CSC) present a subset of cancer cells which is thought to be responsible for tumor growth. That is why CSC are also named “tumor initiation cells”. Additionally, it was shown that CSC are resistant to chemo- and radio-therapies which suggests that these cells can be responsible for tumor relapses after these treatments. Experimental data in cancer cell lines have shown the intriguing *phenomena of CSC population stability*, which means that isolated CSC population rapidly stabilizes at its characteristic level (the relative proportion of CSC in a whole cancer population). We suggest a mathematical model of cancer cell population dynamics, based on experimentally measured dynamics of CSC population stabilization and including main parameters of cell population growth.

We have computationally predicted probability of different scenarios of cancer cell behavior for each experimental case with measurable growth parameters. Moreover, we provide an analytical tool for elucidating important biochemical factors responsible for a particular dynamics of CSC population.

The results may have important implications in therapeutic, because the destroying of a set of factors underlying CSC stability may help to avoid tumor relapses.

## Introduction

Stem cells are undifferentiated cells present in very low numbers in most of the tissues. These cells are responsible for tissue renewal and homeostasis, by giving rise to non-stem cells that proliferate and further differentiate in specialized cells.. Stem cells show very specific features, notably concerning cell division: they are able to undergo asymmetrical division, dividing into a stem cell and non-stem cell; also, the rate of stem cells division is very low as compared to that of non-stem cells.

It has been demonstrated that in most malignant tumors, cancer cell populations appear to include a rare stem cell-like subpopulation believed to be responsible for the initiation and maintenance of the tumor in animals [1–7]. This subpopulation can be detected and purified using specific cellular probes or cell surface markers. *In vitro*, purified cancer stem cells (CSCs) are able to reconstitute the population heterogeneity whereas, in contrast, purified non-stem cells can not. Also, CSCs were shown to be highly tumorigenic in xenografts experiments, and to be responsible for cancer metastasis, i. e. colonization of various tissues by the primary tumor. Because of these features, cancer stem cells are also called tumor-initiating cells [4].

However, it is clear that cancer stem cells do not always comprise all the features of normal stem cells. For example, CSCs may have a diminished capacity to undergo asymmetrical division compared to normal stem cells [3,1–10].

CSCs existence has now been demonstrated in most of solid and hematologic tumors [5,7,9,11–13], with very well described common functional features, which are their indefinite self-renewal capability, existence of asymmetric divisions, and also the resistance to chemo- and radio-therapies [1,2,6,7,14]. The last one suggests that CSCs can be responsible for tumor relapses following chemo- or radio-therapy, which may have important implications in therapeutic, as most of the current treatment targets a regression of the tumors mass without accounting for the tumor functional heterogeneity.

Various mathematical models have been proposed for describing the dynamics of stem cell populations, both normal [15,16, and references there in] and cancer [17,19–21]. These works suggest two different concepts for description of stem cells population behavior. One concept is based on the principle that stem cells act according to their intrinsic program, which may be deterministic or stochastic [20–22]. Alternatively, a concept of self-organization of stem cells [23–25] suggests modeling of the entire system of cell-cell and cell-environment interactions, for which some authors also consider a stochastic behavior. The analysis of available experimental data is not decisive to validate any of these models, but seems to be in contrast with simple models which do not include communications between cells.

However, the possibility of dedifferentiation of non-stem cancer cell to stem cell is one of the most important and yet unsolved question about CSC behavior, which has been, so far, addressed in only few of these works [19,21]. These (stochastic) models base their principle on the assumption that all possible transitions in subpopulations can occur spontaneously, with some probabilities.

Also, only two modeling approaches were suggested for gaining insight into the intriguing *phenomena of cancer stem cells population stability*. Experimental data in cancer cell lines harboring detectable cells with CSC features have shown that over several years of cell passage the relative amount of cancer stem cells fluctuates around a basal level, characteristic for each particular cell line (Figure 1, dotted red curve). Moreover, it has been shown that isolated cancer stem cells, can rapidly recapitulate the heterogeneity of the parental cell line, while, coincidently, the relative proportion of cancer stem cells stabilizes at its characteristic level (Figure 1, dark blue curve).

**Figure 1.**
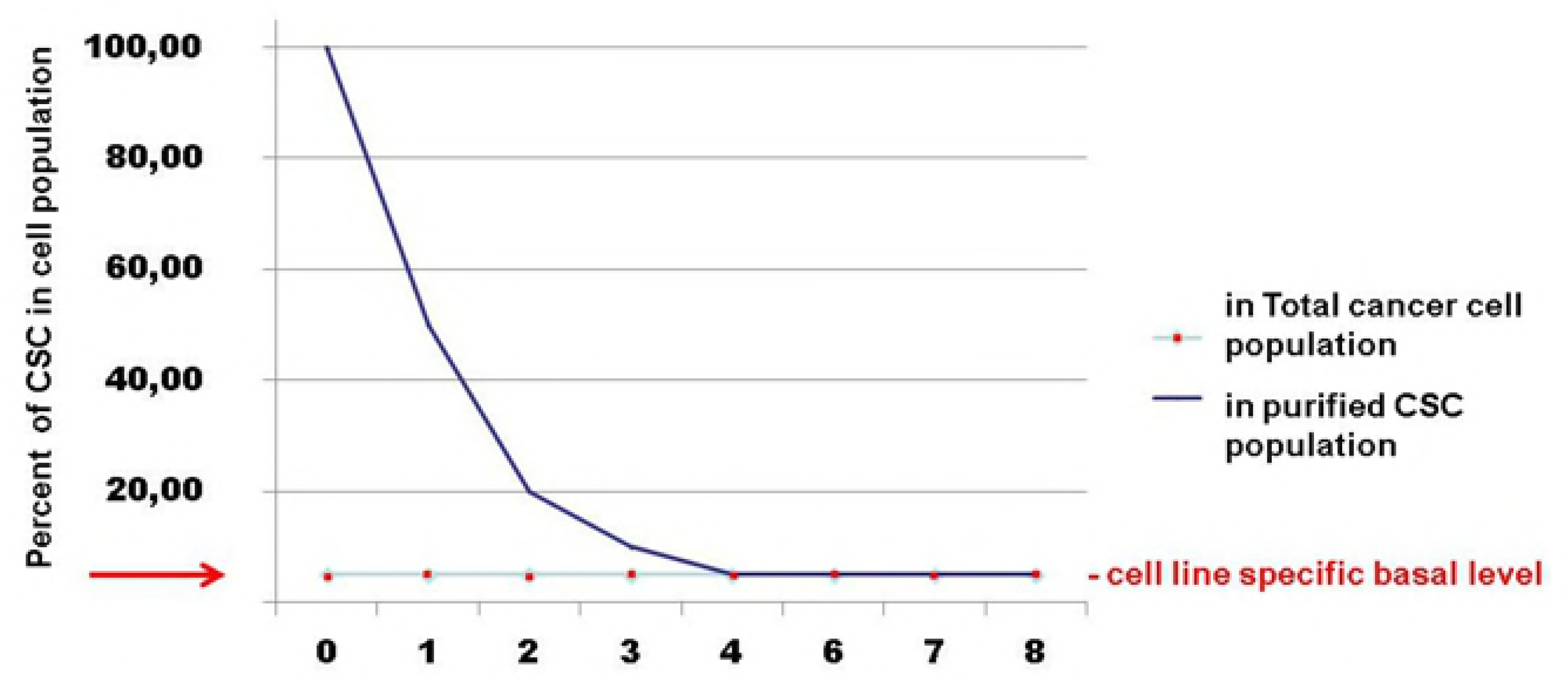
Stabilization of Cancer Stem Cells population in cell culture shown in several cancer cell lines in vitro. The actual curve, showing a percentage of CSC over time, is a generalization of experimental data obtained for human lung adenocarcinoma cells (A549, data provided by A.Harel-Bellan) and for breast cancer cells (SUM159, data provided by C.Ginestier). Human lung adenocarcinoma cells and breast cancer cells were FACS-sorted using CD133 antibodies or ALDEFLUOR assay, respectively. CD133 or ALDEFLUOR positive cells were immediately cultured in their respective culture medium, and analyzed using cell-line specific CSC probes every day thereafter.

One work discussing this phenomenon models the CSC behavior as a Markov process [19]. The model is based on a concept of stochasticity in single-cell behaviors and does not consider the factor(s) of cell-cell communications. Hence, this approach does not allow taking into account the factors involved in regulation of this behavior, and thus, does not address other hot questions in cancer biology, tightly connected to this regulation.

In our previous work [26,27] we have constructed and analyzed the mathematical model which took into account this intriguing characteristic of CSC population behavior. We have suggested a concept of an instructive role of cell-to-cell signaling influencing the parameters of the cell behavior and therefore leading to CSC population equilibrium.

The constructed mathematical model accounts for every possible scenario of cell behavior determining cancer stem and non-stem cells fates, i.e., types of cell division (symmetric or asymmetric), direct transition (differentiation or dedifferentiation), cell death. The analysis of the model helped to elucidate some important characteristics of cancer stem cells evolution, in particular, a set of parameters of cell growth, indicating the necessity of non-stem to stem cell transition for each concrete case of measured cell population kinetics.

In this work we expand this mathematical model to address the question of “instructive signal(s)” underlying the phenomena of cancer cell population stability, aiming to provide meaningful predictions on its dynamics and nature.

We demonstrate that using experimental data of cancer stem and non-stem cells population kinetics measured in the context of CSC population stabilization, our model is able to infer corridors of time-varying probabilities of cancer cell fates that provide significant insights into the cellular dynamics of heterogeneous tumors. Next we show how the set of curves of probabilities of each cell fate scenario can help to identify a set and kinetics of secreted factors responsible for cells population behavior.

We believe that our model can be a useful tool as for resolving important biological questions regarding cancer cell behavior, so for practical medical applications. For example, it may be used to model and analyze the cancer cells population behavior after the chemotherapy treatments, resulting in a perturbation of the total population structure, which might be specific for each concrete tumor cell type.

## Results

### 1 Mathematical Model and Previous Results

In our previous work [26] we have suggested a model accounting for all possible scenarios of cell behavior (cell divisions and direct transition) for stem and non-stem cells, and assumed that each scenario can occur with some probability (Table 1):

**Table 1.** Possible scenarios of cell behavior.

| Stem cell(S) |  | Mature cell (D) |  |
| --- | --- | --- | --- |
| scenarios | probability | scenarios | probability |
| $S \rightarrow S + D$ | $p_1$ | $D \rightarrow D + D$ | $q_1$ |
| $S \rightarrow D + D$ | $p_2$ | $D \rightarrow S + D$ | $q_2$ |
| $S \rightarrow S + S$ | $p_3$ | $D \rightarrow S + S$ | $q_3$ |
| $S \rightarrow D$ | $p_4$ | | |

Though we include the mode 3 for mature cells (giving two S cells) in a Table 1 as theoretically possible, in a model we will assume that its probability *q*_3_ =0, as it is less relevant biologically. The probabilities of scenarios for stem cells (S) and mature cells D should satisfy the usual restrictions on probabilities:

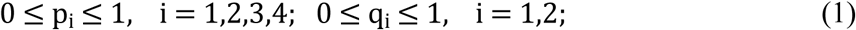

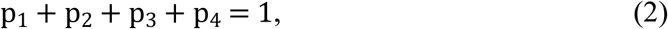

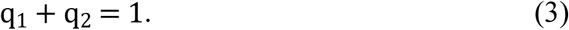

We considered that cell divisions/transition occur with the rate λ_1_ in stem cells (S) and with the rate λ_2_ in mature cells (D), and that death rates are γ_1_ and γ_2_ in S and D cells respectively.

We need to comment that though it seems more plausible to consider that each mode of cell divisions or transition occurs with its own rate (which can be constant or variable with time), the available experimental data indicate that there is no measurable difference in the rate of cell divisions within the sets of stem or non-stem cells, independently from a scenario (C.Ginestier, unpublished data). Thus we consider in our model one constant rate of cell divisions (or direct transition) for stem cells, and another constant rate of cell divisions for non-stem cells.

The proposition (Table 1) gives a system of differential equations for dynamics of *S* nd *D* cancer cells:

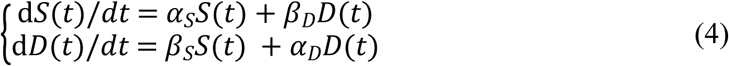

with coefficients, depending on probabilities of scenarios p_i_ and q_i_, and on parameters λ_1_, λ_2_, γ_1_, γ_2_ (growth and death rates):

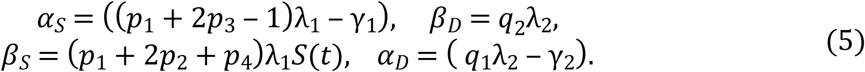

The system can be re-written as:

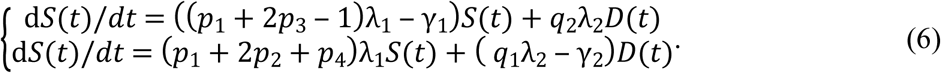

In our previous work, using the system (6), we have analyzed the time-dependent evolution and the asymptotic behavior of the percentage of cancer stem cells *s*(*t*) in a cancer cell population, as illustrated in the experimental curve presented in Figure 1:

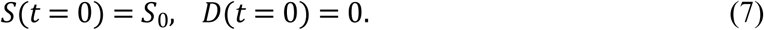

The percentage of stem cells by definition is:

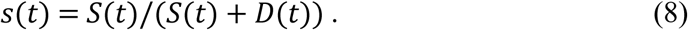

The analysis of the model has given the answers to important biological questions:

1. Can a cell line-specific equilibrium level of CSCs be explained only by intrinsic characteristics (i.e., growth parameters) of a cell line?
2. What are the necessary and sufficient conditions for the stabilization of Cancer Stem Cells population?

The most important conclusions which we have obtained can be briefly summarized as follows.

1. Is it not possible to achieve CSC stabilization with only standard (considered as predominant) modes of S and D cell divisions, namely with only asymmetric division for S cells, and symmetric divisions, producing two D cells, for D cells.
2. Assuming the existence of different modes of both S and D cells behavior (divisions and direct transitions), and considering only a set of *constant* parameters rate, a stabilization phenomena can be achieved only in one of two cases:

a. in admission of the *D*→*S* transition scenario, i.e., asymmetric division of *D* cells (*q*_2_ > 0);
b. without *D* to *S* transition, only if *S* cells have very high frequency (probability) of symmetric divisions into two *S* cells (*p*_3_), and if the set of growth and death rate parameters λ_i_,γ_i_, satisfies the condition:

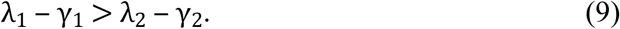

The last one looks to be contrary to the existing experimental data, because when γ_1_ = γ_2_ it gives λ_1_ > λ_2_, while a condition reflecting a relatively slow speed of stem cell divisions means λ_1_ < λ_2_; however, as we have already mentioned, this can be different in cancer stem cells versus normal stem cells, and especially in the abnormal cases of perturbed (purified) cancer cells population.

### 2 Corridors of probabilities of cell fate scenarios

In the presented work we continue analysis of the model aiming to solve the following problems:

determination of time-varying corridors of probabilities of different cell fates, given the dynamics of cancer cells populations;
determination of a cell-to-cell communication factors, influencing time-varying probabilities of cell behavior (division, direct transition) scenarios.

First we address the *Question 1*:

W*hat are the necessary conditions on parameters λ_i_,γ_i_, *p*_i_,*q*_i_ a model (6) for achieving a complete fitting of an experimentally measured curve of the percentage of CSC over time displaying a stabilization phenomenon after CSC perturbation?*

For this search we have used experimental data, provided by A.Harel-Bellan and C.Ginestier, shown on the Figure 1, blue curve, namely the percentage of CSC at each time point starting from purified to 100% CSC population and up to the time of its equilibrium (stable state), achieving at 1-5% depending on a particular type of cancer and cell line.

The percentage of CSC *s*(*t*) was calculated by Fluorescence Activated Cell Sorting (FACS) analysis measuring amounts of S and D cells at each time point of a kinetics over eight days, and presented as in (8).

The results of our analysis and numerical simulations show that a good fitting of model (6) to the experimentally measured curve s(t) (determined by the least squares method, with an average deviation equal ~ 5% or less) can be achieved *if and only if the set of parameters λ_i_,γ_i_, *p*_i_,*q*_i_ changes over time*.

Due to the experimental data, discussed in the Section 1, we may admit for the next analysis that parameters changing with time should be the probabilities of scenarios *p*_i_(*t*),*q*_i_(*t*) while parameters λ_i_,γ_i_, may be accepted to be constant.

This leads us to the *Question 2*:

*Given the dynamics of percentage of CSC s(t), is it possible to find functions* p_i_(*t*), i = 1,2,3,4, q_i_(*t*), *i* = 1,2 *for a given constant set of* λ_i_ *and* γ_i_ *?*

In order to solve this problem we first considered the following *Hypothesis: all changes in cell behavior scenarios up to stabilization should be minimized (the sum of all changes of p*_i_(*t*) *and q*_i_(*t*) *should be the minimal possible ones)*.

This means that the function v:

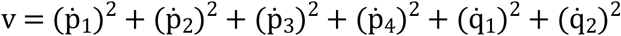

where 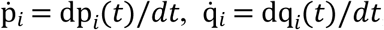, should be minimized, i.e.:

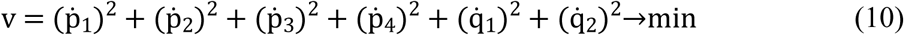

Next we rewrite the system (6) and the conditions (2) and (3) as:

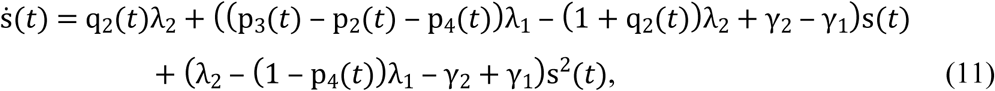

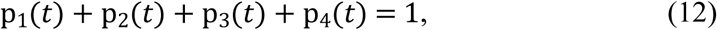

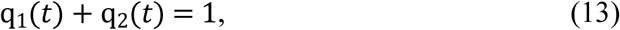

where 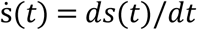.

This gives us a system:

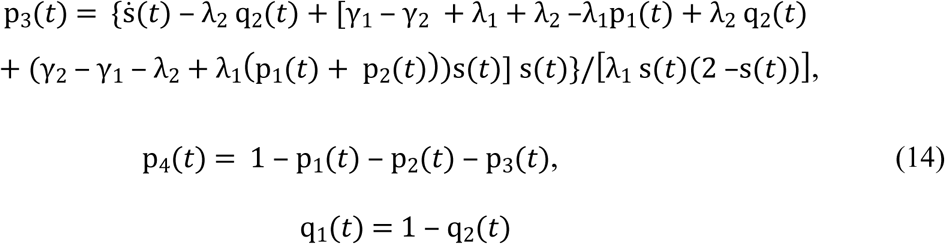

from which we can conclude that the function v depends only upon three variables, they are (for example) p_1_(*t*),p_2_(*t*),q_2_(*t*). Thus, for minimizing it we need to solve 3 equations:

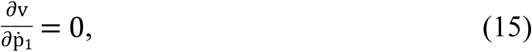

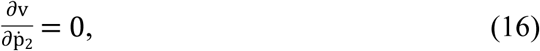

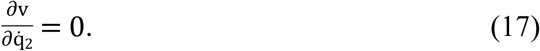

Thus, for six variables p_i_(*t*) and q_i_(*t*) we have a system of 6 equations (11), (12), (13), (15), (16) and (17). For the solution of that system we have to add three initial conditions:

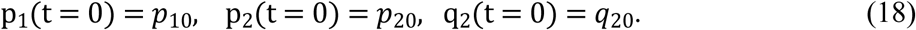

All solutions for each p_i_(*t*), q_i_(*t*) corresponding to all possible sets of initial conditions (18), which satisfy the condition (1) for all time period T will provide a set of six corridors (ranges) of probabilities of scenarios p_i_(*t*), q_i_(*t*) of cell behavior in a given experimental case.

This means that *given measured s(t) and constant set of* λ_i_,γ_i_*, we can determine corridors of all possible probabilities of scenarios* p_i_(*t*), q_i_(*t*) *varying with time*.

The results of the calculations of all possible corridors for the different sets of biologically relevant parameters λ_1_,λ_2_,γ_1_,γ_2_ are presented on Figure 2 (the program code can be found at https://cloud.mail.ru/public/J2jk/637e7Bpcr). The considered rates of non-stem cell division (λ_2_) were taken from experimental measurements in different cancer cell lines, while the rate of stem cell division (λ_1_), which is difficult to measure, was chosen in order to vary the ratio of stem/non-stem division rates (1:1; 1:5; 1:10), corresponding to a statement about slower rate of stem cells divisions. The considered variants of cell death rates (0.1; 0.5) were taken from experimental measurements.

**Figure 2.**
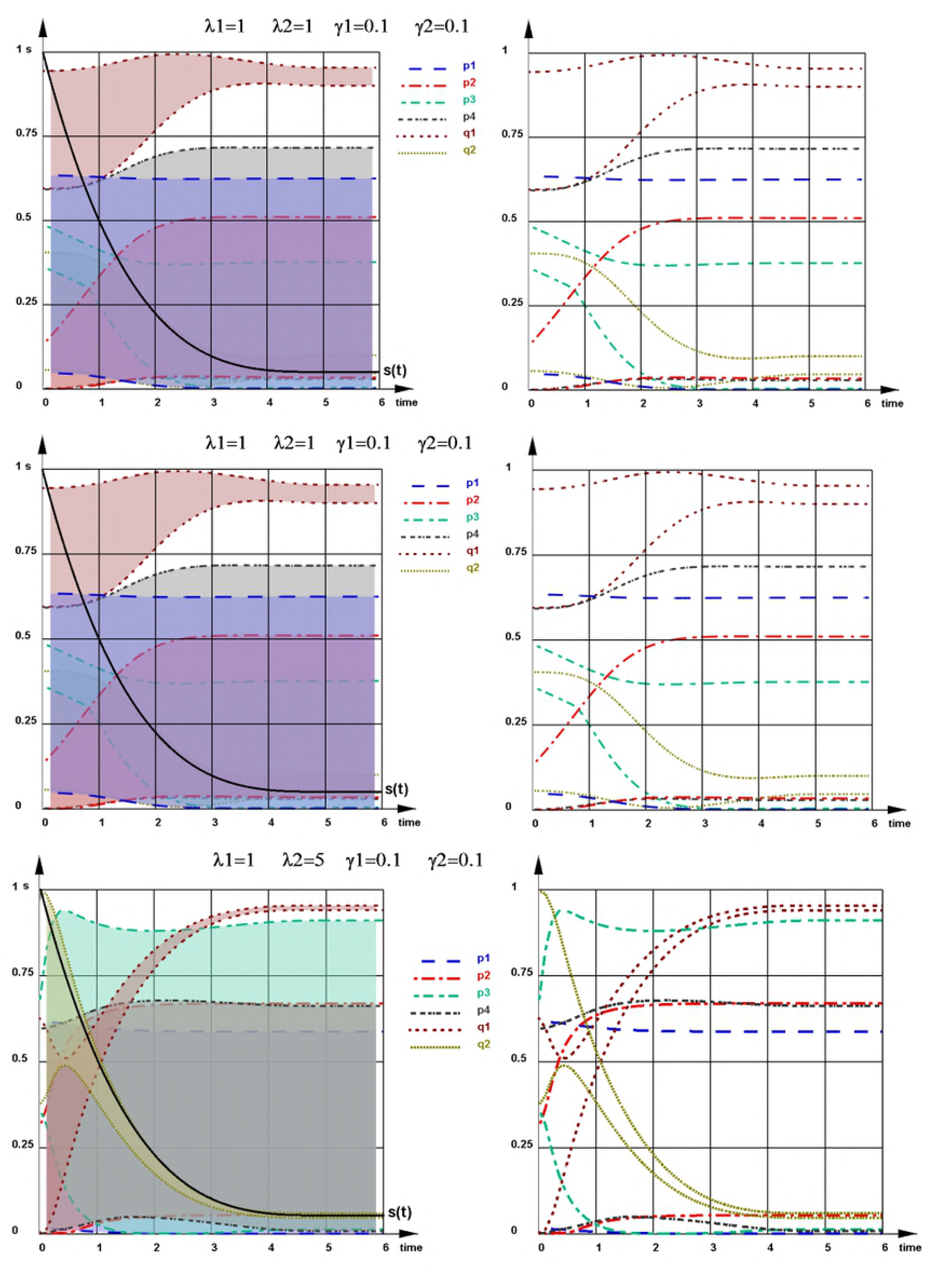
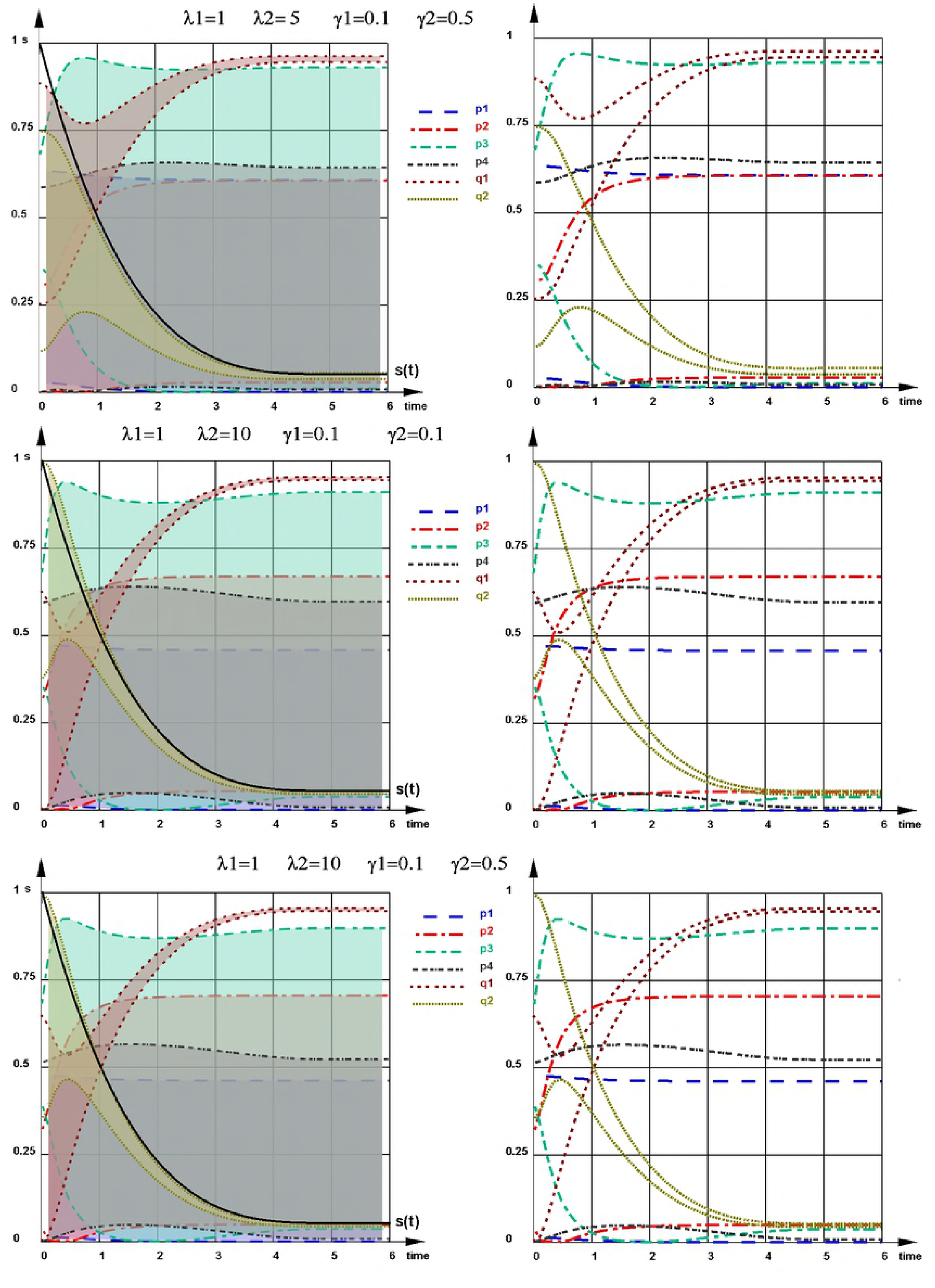
The ranges (corridors) of all possible solutions for probabilities p_i_, q_i_ for the given experimental curve of CSC sub-population dynamics (s(*t*), black curve). The boundary of corridors are determined by satisfying the condition (1). Left panels: each “corridor of probability” is shown in sheeted color. The experimental curve of CSC sub-population dynamics s(*t*), is shown in light green color. Right panels: “corridor of probability” is marked by max and min colored lines for the following sets of λ_1_,λ_2_, γ_1_, γ_2_.

It is important to note, that the boundaries of each corridor can be (and usually are) comprised from the parts of different functions p_i_(*t*) (or q_i_(*t*)) existing inside a particular corridor.

However, these boundaries can help to evaluate the possibility or impossibility of a questionable scenario(s) for each concrete case (in a particular experimental system).

For example, this simulation gives an answer to the question of the *necessity of non-stem to stem cell transition scenario* in a course of CSC population stabilization and next maintaining of an equilibrium level. As it can be seen from the Figure 2, for all considered cases of a biologically relevant set of cell division and cell death rates λ_1_,λ_2_, γ_1_, γ_2_, the lowest boundary of the corridor of probability q_2_(*t*) (corresponding to non-stem to stem cell transition) appears to be higher than zero, at all or at least at some time points.

Another important question is about the existence of other types of stem cell behavior scenarios except the asymmetrical one, which is postulated to be predominant by current cancer biology. The obtained results showed that for considered biologically relevant sets of parameters λ_1_,λ_2_, γ_1_, γ_2_ the highest level of possible probability p_i_(*t*) (the upper boundary of its corridor) is around 60%, being much lower for some cases (Figure 2).

These important conclusions are illustrated more detailed on the Figures related to the next question been considered. For many practical applications it is very important to be able to find a unique solution for each probability p_i_(*t*), q_i_(*t*). For that end we have to address the *Question 3:*

*Given a measured curve s*(*t*)*, and parameters* λ_i_,γ_i_*, what additional data are necessary and sufficient for getting a unique solution for each* p_i_(*t*) and q_i_ (*t*) *?*

From the analysis done for the question 2 it is clear that additional data which are necessary for getting a set of unique functions p_i_(*t*) and q_i_ (*t*), is a set of three initial conditions (18).

We have explored this possibility and present two examples of the determination of such a set of unique functions p_i_(*t*) and q_i_ (*t*) on the Figure 3 and 4. Six different cases of given parameters λ_i_,γ_i_, (the same as on the Figure 2) are considered. In the first simulation (Figure 3) we chose the initial conditions (18) in way providing *q*_20_ be the minimal one. For the other simulation we set the initial conditions demanding *p*_10_ to be the maximal one (Figure 4).

**Figure 3.**
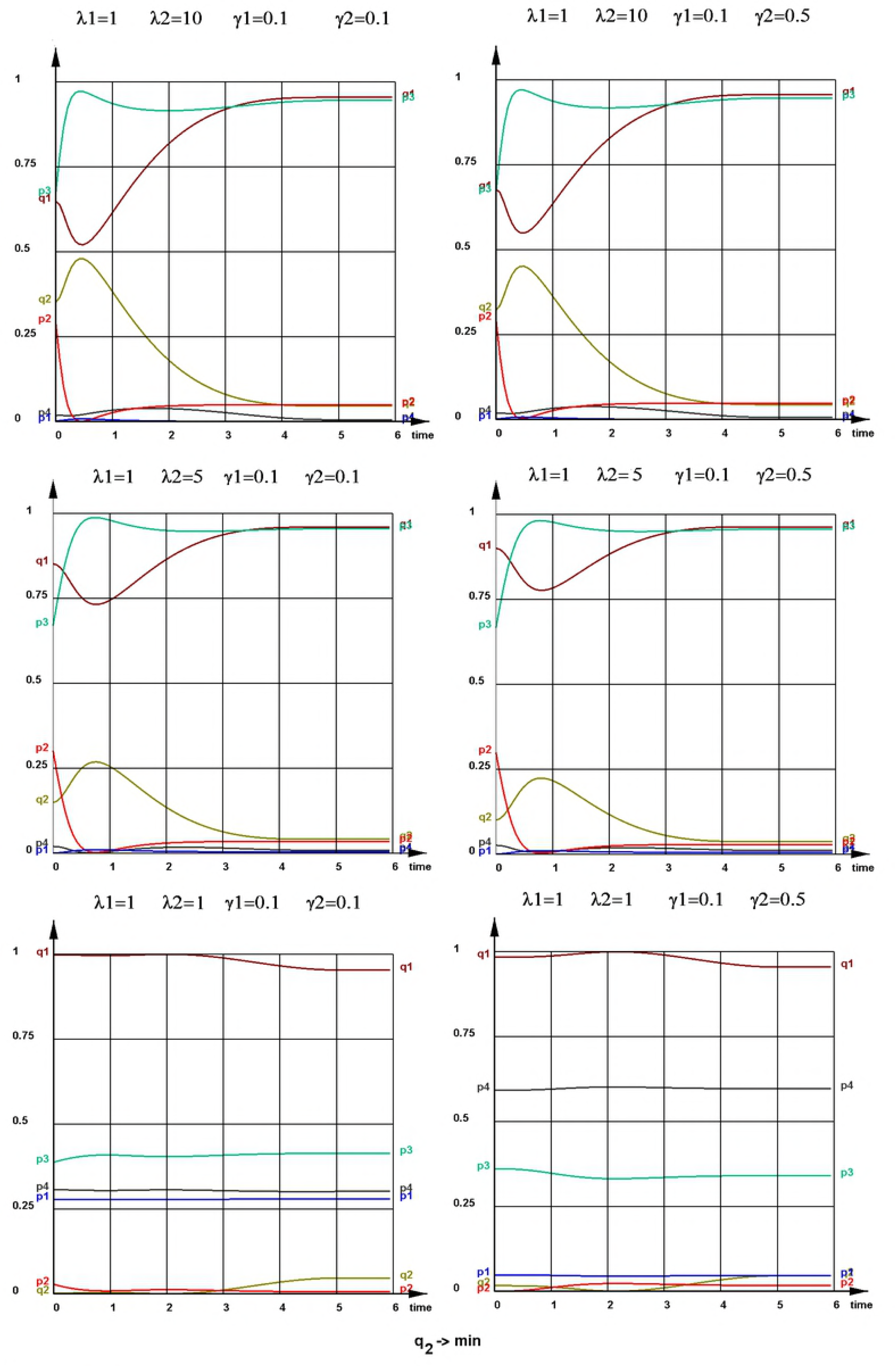
The set of p_i_(*t*) and q_i_(*t*) functions is uniquely determined by the choice of initial conditions requesting *q*_20_→*min*.

**Figure 4.**
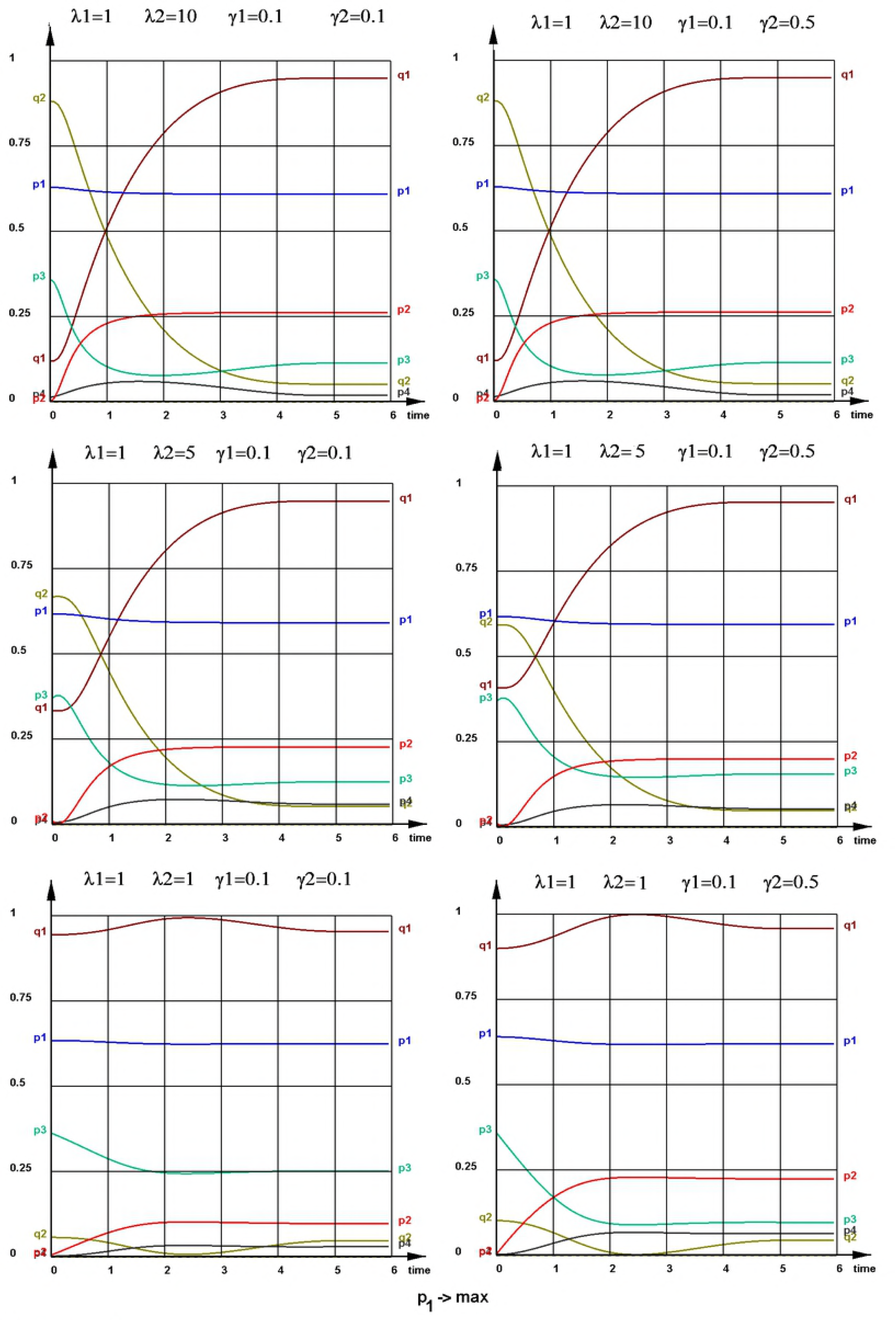
The set of p_i_(*t*) and q_i_(*t*) functions is uniquely determined by the choice of initial conditions requesting *p*_10_ →*max*.

### 3 Determination of underlying field factors, responsible for the time-varying cells behavior

In our previous work [26] we have suggested that the coordinated dynamical change of the parameters of cell behavior, resulting in the phenomena of cancer stem cells subpopulation equilibrium, occurs in response to a multiparametric biochemical signal produced in a system. In the simplest case, it can be a set of secreted factors in the media influencing cell behavior. We notated this signal as an *underlying field* u(*t*), where u(*t*) generally speaking, is a matrix of factors u_ij_(*t*), where each factor *i* may be produced by S or D cell at some time point t and have an influence *j* on a behavior of S or/and D cell, possibly with some time delay t+r.

Here we further explore this idea, assuming that a field of secreted factors influences probabilities of cell fate scenarios, and thus a structure of each corridor of probabilities p_i_ and q_i_ depends on a changing set of concentrations of secreted biochemical compounds.

This means that in the equation (10) all probabilities of division scenarios of S and D cells (p_i_(*t*) and q_i_(*t*)) are the functions of an underling field u(*t*), changing with time:

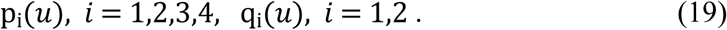

We can also consider possible a dependence of each factor u*_k_*(*t*) on the amount of cells S and/or D, due to its production by one of these type of cells correspondingly.

The task of identification of molecular factors involved in the underlying field formation raises the *Question 4:*

*Given a set of unique functions of probabilities *p*_i_ and q*_i_ *is it possible to find a set of factors* u_*k*_(*t*) *responsible for their dynamics?*

In order to determine the factors u*_k_*(*t*) which influence the evolution of probabilities p_i_(*t*) and q_i_(*t*), we will perform decomposition of given functions and q_i_(*t*) over the functions u*_k_*(*t*).

We will use the following form for the function u(t):

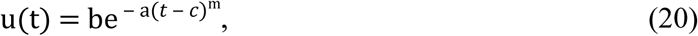

where a is the coefficient reflecting the width of a particular curve u(*t*), b is its height, c is the position of the curve on the time axes, m is the shape (sharpness) of the curve (Figure 5).

This form, by varying the coefficients *a,b,c,m* allows to model many different kinetics of factors, e.g., linear ones with different parameters, constant ones, exponential, trapezoid, etc. The test calculations and numerical simulation have shown that it is possible to set the value of the coefficient m = 6 for all factors, while the coefficients a, b, c should be selected by least-squares fitting.

**Figure 5.**
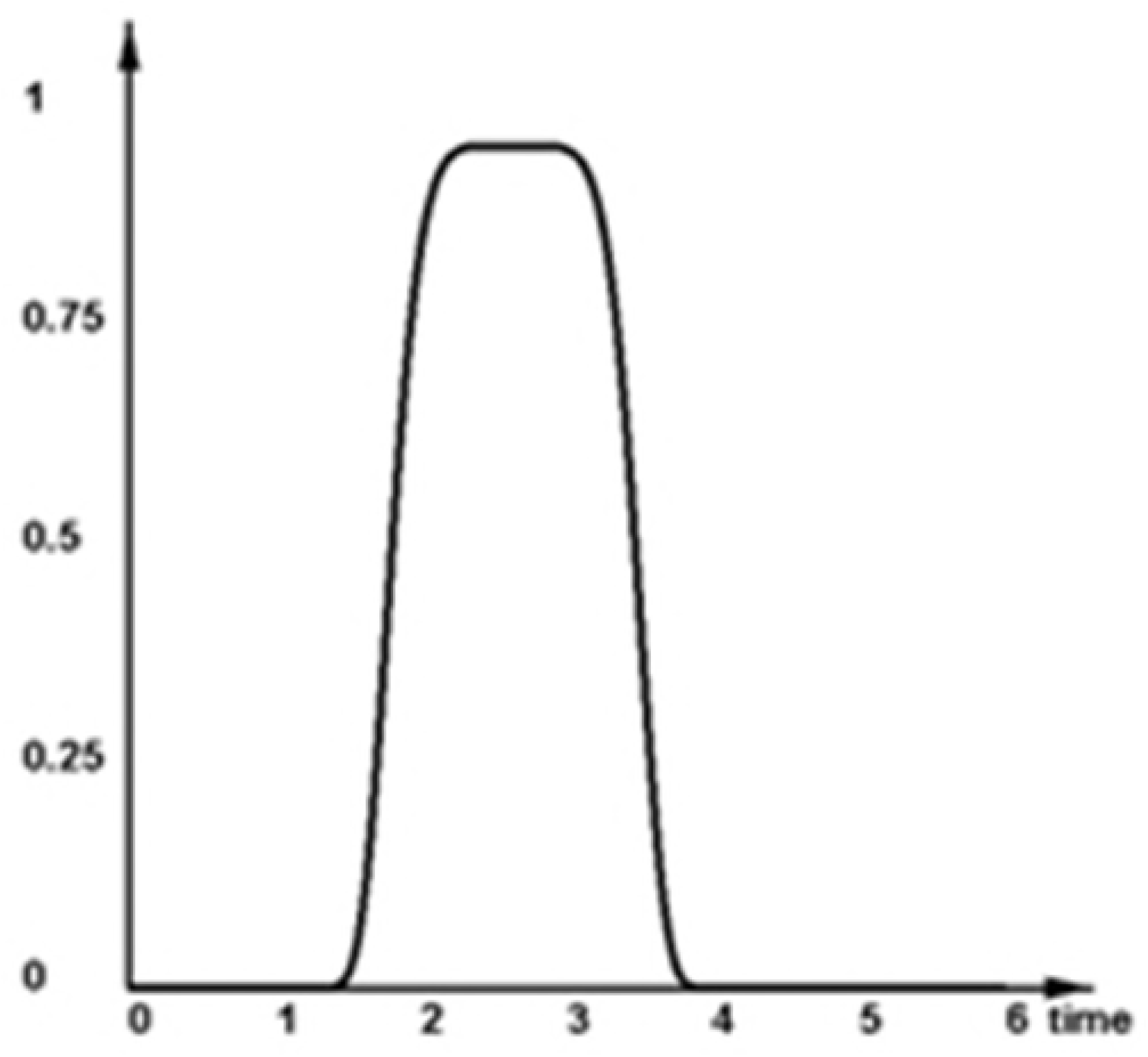
The function u(t) for approximation the parameters and behavior of factors of influence.

Next for the sake of generalization, we will consider one variable *y*_i_, *i* = 1…6, for all probabilities six probabilities *p*_1_,*p*_2_,*p*_3_,*p*_4_,*q*_1_,*q*_2_.

Thus, the probabilities *y*_i_(*t*_*n*_), *i* = 1,…,6
will be:

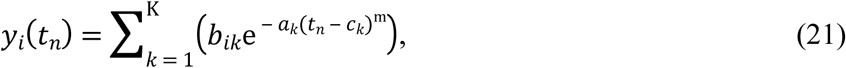

where K is the number of u(t) factors considered, n = 1,…N is the experimental time-points Using the least squares method, we will find the coefficients *b*_*ik*_,*a*_*k*_,*c*_*k* in the_ expression (21) such that they provide a minimum of the total quadratic deviation:

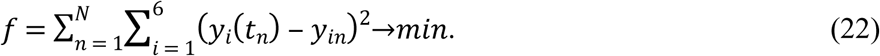

For that it is necessary to ensure equality to zero of the expressions:

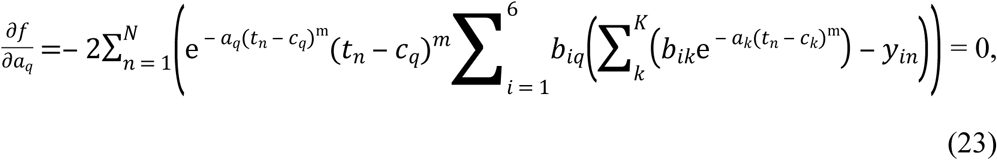

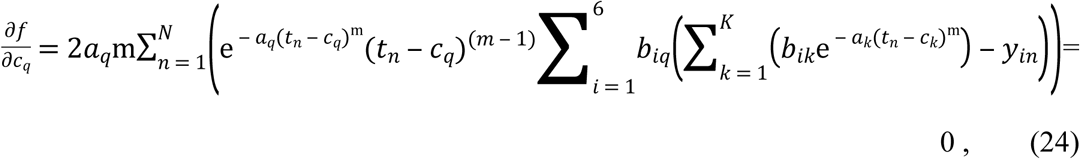

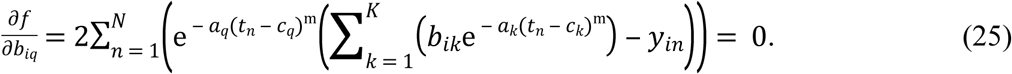

By this we obtain a system of equations (23-25) for determining *b*_*ik*_,*a*_*k*_,*c*_*k*_, *k* = 1…6, *i* = 1…6, which was solved by the gradient descent method, starting with test initial values and obtaining the minimum of the function (22). The number of factors k is determined from the condition that the mean deviation exceed the given value of permissible variation ε:

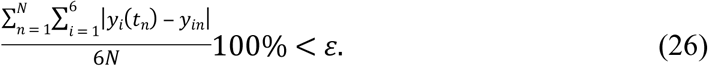

The search starts from suggested initial number of factors K, and in the case when condition (26) is not fulfilled for a given ε, should be repeated for K+1, and further on until the satisfaction of (26).

Next a corresponding computer program, automatically performing all necessary computations for a given ε, was developed and applied for the particular set of curves *p*_1_(*t*),*p*_2_(*t*),*p*_3_(*t*),*p*_4_(*t*),*q*_1_(*t*),*q*_2_(*t*) presented on the Figure 3 (the program code can be found at https://cloud.mail.ru/public/J2jk/637e7Bpcr). The results of computations, determining the number and pattern of u*_k_*(*t*) factors, providing the satisfaction of the condition ε ≤ 1% for each case (the same sets of λ_i_,γ_i_ considered before), are presented on the Figure 6, where the height of each factor (which is proportional to the normalized concentration of putative secreted factor in a media) is shown as

**Figure 6.**
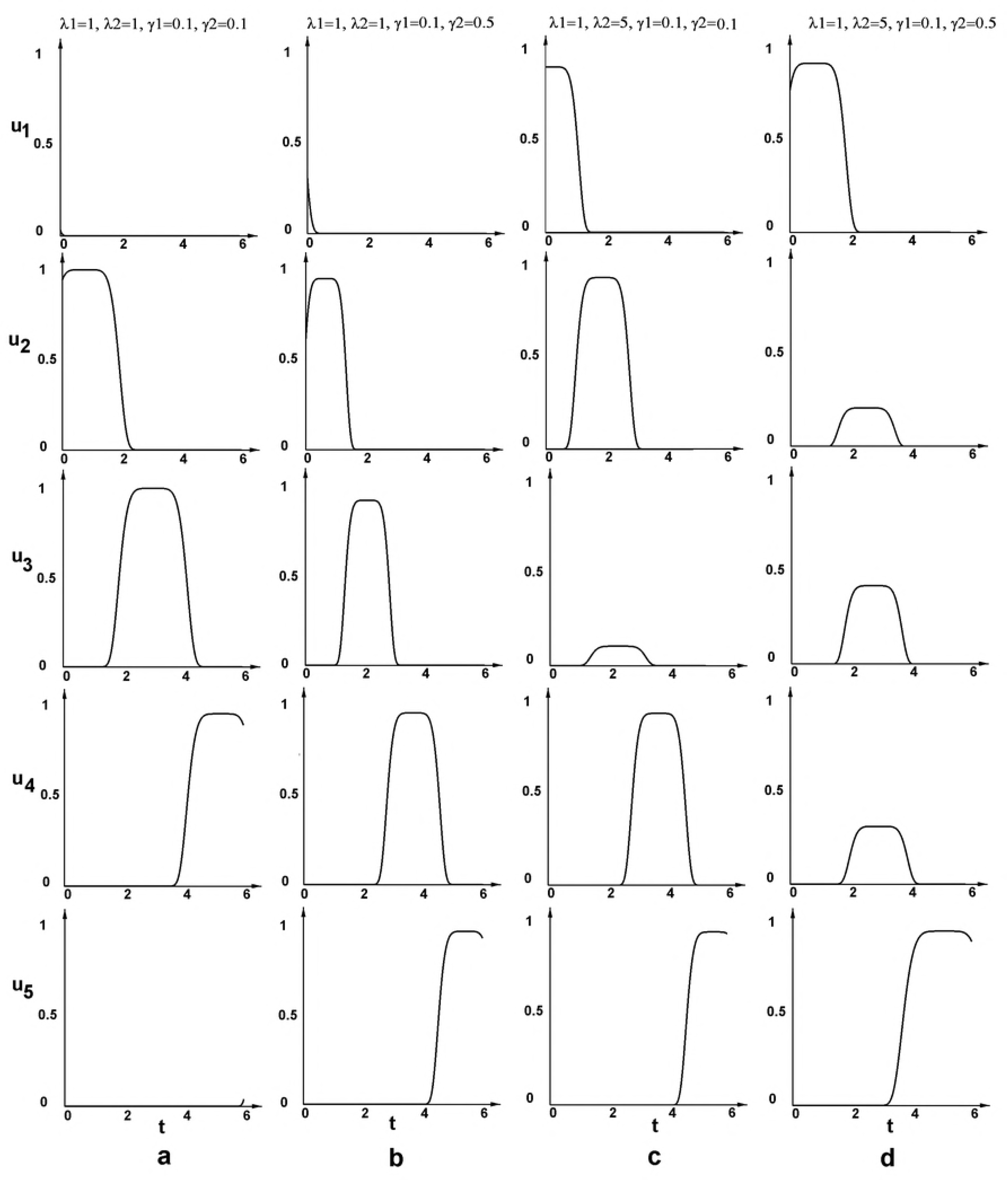
Underlying field u*_k_*(*t*) functions for four considered sets of λ_i_,γ_i_.

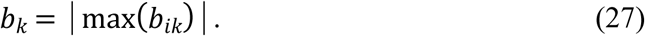

The search started from suggested initial number of factors *K* = 4, and has elucidated the most plausible number of major u*_k_*(*t*) factors for each experimental case (which varied from 3 to 6 ones depending on a case), together with their specific patterns (shape, height and timing).

It was found that in 3 cases from 6 explored, the starting number of factors *K* = 4 was enough to determined a set of essential influencing field factors u*_k_*(*t*) with sufficiently good value of variation ε.

Moreover, it two cases, namely, for the set λ_1_ = 1,λ_2_ = 1,γ_1_ = 0,1,γ_2_ = 0,1 (ε=0,49%; Figure.6a) and for the set λ_1_ = 1,λ_2_ = 10,γ_1_ = 0,1,γ_2_ = 0,1 (ε=1%; not shown), the program has detected only 3 factors as the essential ones, thus showing the capability to find the minimal possible set of factors u*_k_*(*t*) for the best fitting independently of a starting number of factors K. In one case 4 factors were determined: λ_1_ = 1,λ_2_ = 1,γ_1_ = 0,1,γ_2_ = 0,5 (ε=0,47%, Figure 6b). For the other 3 cases, for which with *K* = 4 unsatisfactory ε value was obtained, the search was continued with grater number of factors *K* up to the good result of fitting. Finally, for two cases the final number of essential influencing factors was 5: λ_1_ = 1,λ_2_ = 5,γ_1_ = 0,1,γ_2_ = 0,1 (ε=0, 82%; Figure 6c) and λ_1_ = 1,λ_2_ = 5,γ_1_ = 0,1,γ_2_ = 0,5 (ε=1,0%; Figure 6d), while in one case, for which K=5 gave ε=1,2, the final number of factors K appeared to be equal to 6: λ_1_ = 1,λ_2_ = 10,γ_1_ = 0,1,γ_2_ = 0,5 (ε=0, 75%; not shown).

We expect that these results *can be used as a tool for identifying molecular factors in the media* comprising an underling field for each particular cell line, and suggest the following experimental strategy.

In a course of measurements of CSC population kinetics (corresponding to a one on Figure 1) at each time point, starting from 100% CSC population and up to its steady state, together with the detection of percentage of CSC subpopulation, the Metabolome (Secretome) profiles of a cultural media at each time point should be obtained. This will identify a set of secreted factors that appear, increase, decrease or disappear at each time point (Figure. 7).

**Figure 7.**
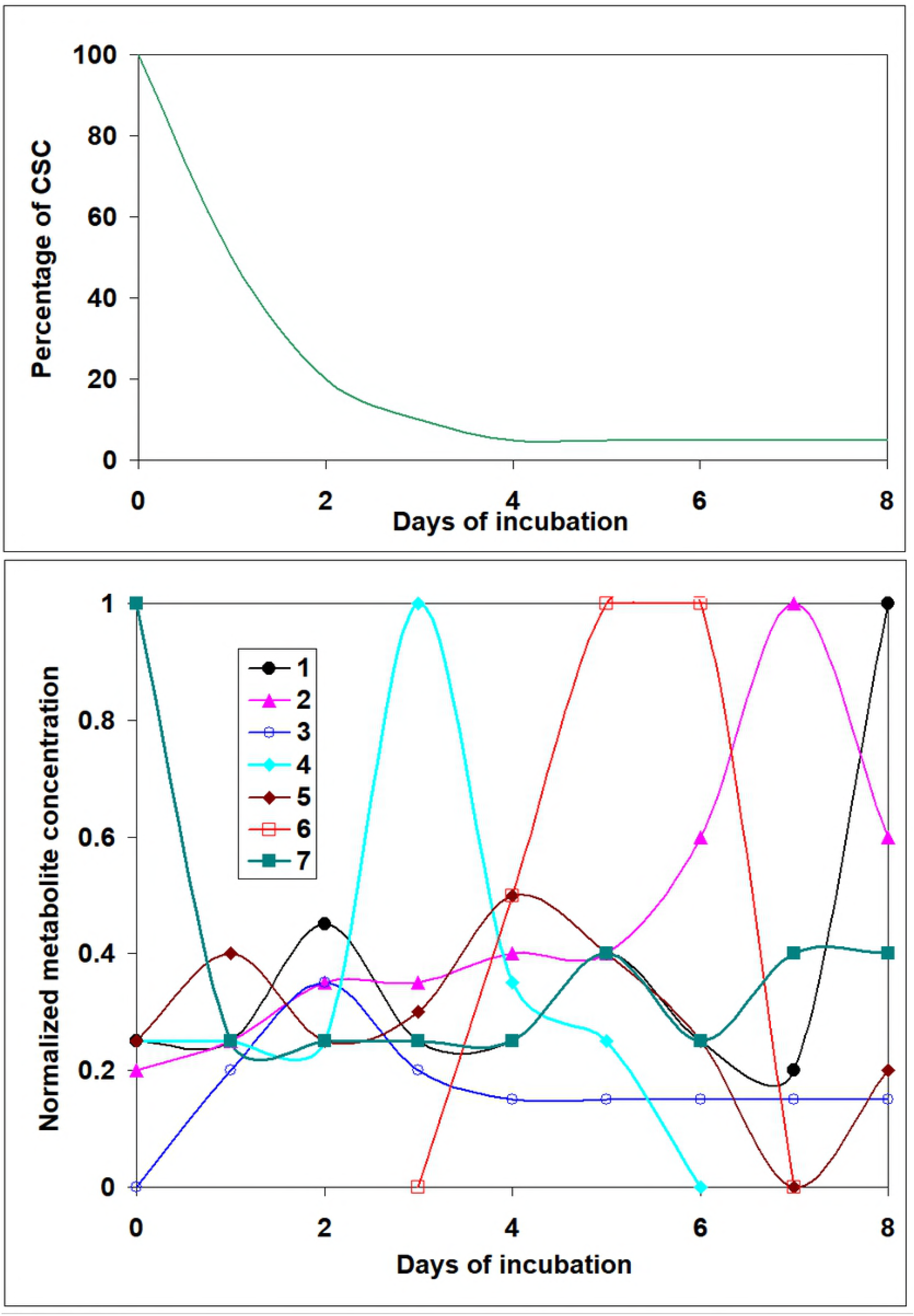
Illustration of Metabolome (Secretome) time-dependent data needed to predict a set of underlying factors involved in the controlling of cell population behavior using a comparison with the results of mathematical modeling.

Next a comparison of the predictions on u*_k_*(*t*) factors obtained by mathematical model for a given case (*s*(*t*), λ_i_, γ_i_) (Figure 6) with the results of the corresponding Secretome profile (Figure 7) should be undertaken. Those biochemical factors, whose timing and pattern coincide with the predicted behavior of the underlying field functions u*_k_*(*t*) will be good candidates for being involved in controlling cell behavior; the height of each factor u*_k_*(*t*) on the Figure 6 corresponds to a normalized concentration of a putative secreted factor in a media. In the descriptive example, presented on the Figure 7, metabolites *N4* and *N6* have kinetics in agreement with the ones found for factors u_3_(*t*) and u_4_(*t*) in Figure 6a.

It is very probable that having several hundreds metabolites in Secretome results, for each u*_k_*(*t*) predicted by the model several metabolites can give a well fitting. In this case both of two variants are possible: a cumulative effect of all (or several) indicated metabolites creates a concrete u*_k_*(*t*) signal, responsible for a changing cell behavior leading to a steady state of CSCs; or only one among them has this special effect. This can be found in a course of next experimental proving of the identified factors being involved in the underlying field formation.

*Important Remark*. Though the system of equations (23), (24), (25) can not be solved uniquely, our computational experiments have shown that by requesting *ε* to be considerably small and by setting the approximation time steps in the program to be 10^-4^ arbitrary units, we get the minimizing of the area of all possible results for each factor u_*k*_(*t*) in such a way that the difference between the various solutions is negligible, because it is below the sensitivity of experimental equipment used for monitoring the factors.

For example, for *ε* ≈ 1% the maximal variation of coefficient b reflecting a concentration of a factor in a media, is 3%, and maximal variations of coefficients a and c reflecting the time of existing of the factor in media, is 3%.

This means that the suggested model can be a used as a tool for determination the exact set of secreted factors in the media, influencing the time-varying cells behavior leading to stabilization of CSC population.

### 4. Predictions of putative production of secreted factors u_k_(t), by S or D cells

The knowledge that factor(s) influencing CSC behavior and responsible for the stabilization of cancer cell population are produced by specific type of cancer cells, namely, by stem or non-stem ones, is extremely important as for biological research, so for possible medical applications. Thus, in order to provide a tool for such a prediction, we assumed that some factors u_k_(t) can be produced mostly (or completely) by S cells, some factors – by D cells, and some – more or less equally by both types of cells.

To account for this possibility, we have considered a function differentiating multiplicative dependence on S or D cells kinetics:

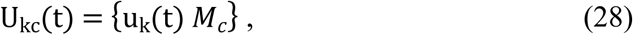

where *M*_c_ = {1,s(t),d(t)}.

This means that having an optimal set of factors u*_k_*(*t*), found at the previous step (Figure.5) for a given set of parameters λ_i_,γ_i_ and given kinetics p_i_(*t*), q_i_(*t*) we will consider all variants of multiplication of each factor u*_k_*(*t*) on each of the functions c = 1 S(t), D(t), with determining *ε* value for each case.

We can assume that the result of differentiating multiplication, which gives the smallest *ε* value for a concrete case, represents the possible dependence of the factors upon the concrete types of cells (*S* or *D* cells). The variant of multiplication on 1 means independence of a factor on *S* or *D* kinetics. The results of the computation U_kc_(t) for two chosen cases are present in the Table 2.

**Table 2.**
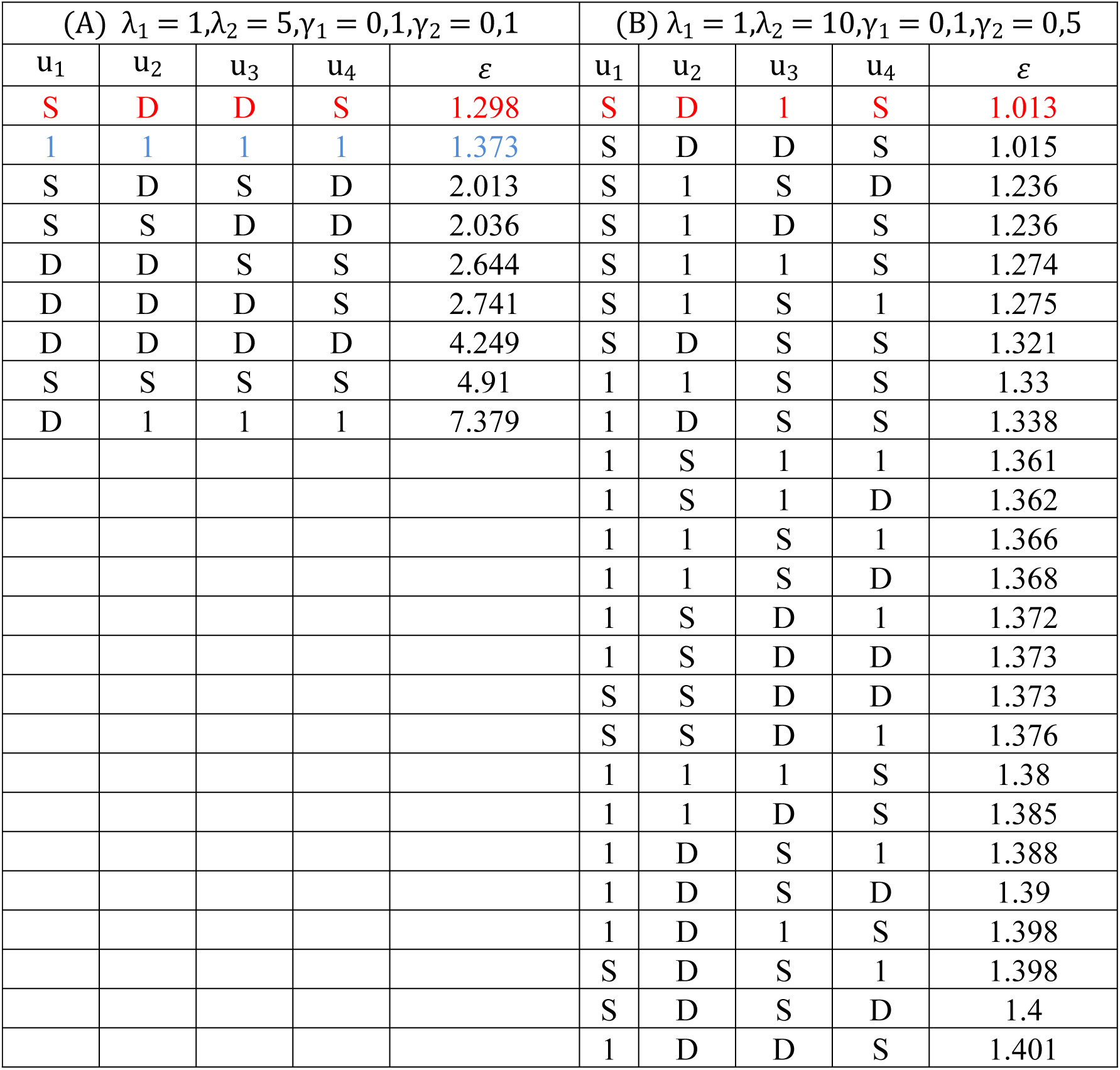
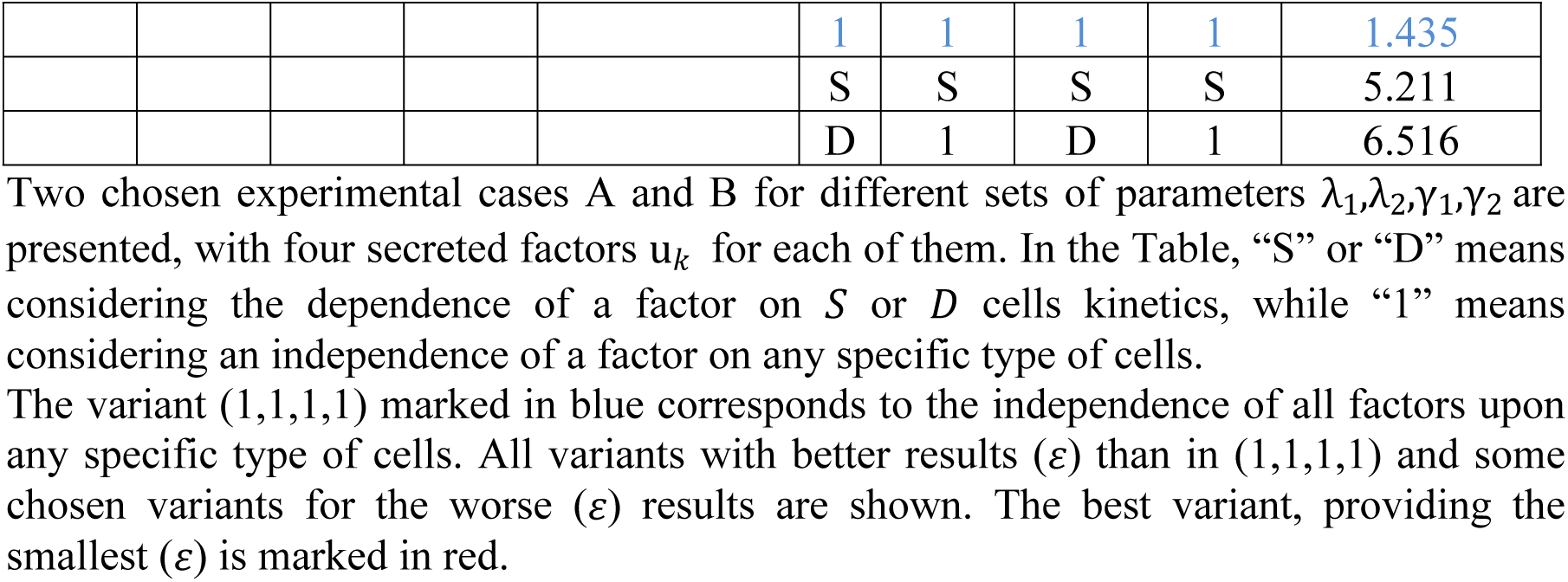
The possible dependence of the factors u_*k*_ upon the concrete type of cells (*S* or *D* cells).

It can be seen that for the case: λ_1_ = 1,λ_2_ = 5,γ_1_ = 0,1,γ_2_ = 0,1, for K = 4 and without differentiating multiplication (corresponding to the variant 1,1,1,1) *ε* = 1.373% (Table 2A). The computation has shown that all other variants of multiplication give worse results (some chosen ones are shown), except for only one variant (*S*,*D*,*D*,*S*), though decreasing the value of *ε* not considerably (*ε* = 1.298%). However, it still can be an indication of differential production of the factors (*t*) by *S* or *D* cells.

Much stronger dependence on *S* and *D* kinetics is demonstrated in the other case (Table 2B): λ_1_ = 1,λ_2_ = 10,γ_1_ = 0,1,γ_2_ = 0,5, where for *K* = 4 and without differentiating multiplication the lowest *ε* value was found as *ε* = 1.435% while in the best variant with differentiating multiplication (*S*,*D*,1,*S*) it decreases up to 1.013. Also, the fact that in this case 25 variants with differentiating multiplication give better results than the variant without considering the dependence on *S* or *D* kinetics (1,1,1,1), can indirectly point on the necessity of concluding this additional calculation in this concrete case. On the other hand, this fact can be an indicator of instability of the found best variant of multiplication. (For this case, all variants with better results (*ε* than in (1,1,1,1) and some chosen variants for the worse (*ε*) results are shown in the Table 2B).

It is important to note, that in order to show the more pronounced difference in multiplicative and non-multiplicative cases, we present in the Table 2 the results of a differentiating multiplication search, which was started from the optimal sets of non-multiplied factors u*_k_*(*t*) obtained with the requirement *ε* ≤ 1.5, which correspond for each of the presented cases to *K* = 4. The application of this search for the best non-multiplicative set obtained in stricter requirement *ε* ≤ 1, which results in 5 factors in the first case and 6 in the second one, gives the same qualitative results, though showing much less relative decrease of *ε*.

This means that in order to find a reliable results about the dependence of the underlying field factors on *S* or *D* cells, one should apply the suggested computational program, starting with the optimal sets of non-multiplied factors u*_k_*(*t*) obtained for possibly high level of *ε*.

## Discussion and Conclusions

The results of our work lead to several important theoretical suggestions, and the basis of which we provide analytical tools to experimentalists for the analysis of cancer cell behavior, which can not be accomplished by biological methods only.

Mathematical modeling of different possible cell fate scenarios suggests that the achievement of cell line-specific equilibrium level of CSC percentage depends on dynamic changes of system parameters, and that these changes are most likely controlled by a field of time-varying cellular interactions. The computational program based on our model allows determining several important specific characteristics of CSC behavior in a given cancer cell population.

First, on the basis of a minimal set of experimental values (rates of cell division, of cell death and CSC population kinetics s(*t*)), our model enables the predictionof cell-line specific modes of *S* and *D* cells divisions, including the unusual *D*→*S* transition. provided that

Second, the model predicts the existence and the dynamics of cell-to-cell interaction factors u_k_(t) influencing the cancer cells behavior (namely, the probability dynamics of cell division modes p_i_(t) and q_i_ (t)).

Finally, the model predicts the dynamic profile of cell-cell interaction factors u_k_(t) specifically produced by S (stem) or D (non-stem) cells in the cancer cell population.

In summary, we provide a mathematical tool using a limited amount of experimental measurements to help experimentalists gain insight into a broad array of cancer cell fates.

While our model’s predictions have still to be experimentally confirmed by a set of biological experiments (e.g. supplementation of a factor in media, or KD by small interfering RNA followed by next detecting the shift in CSC kinetics), we could anticipate the development several practical fundamental and practical applications.

In the future, the proposed work may give a tool for the evaluation of of the risks of cancer to relapse upon radio- or chemotherapy, based on the experimental measurement of its CSC kinetics of. Also, it may help find the proper treatment achieving the most efficient suppression of each cancer subpopulations growth, together with the proper treatment schedule to achieve the best therapeutic result by considering the predicted *underlying field* behavior.

Next the analysis of the experimental kinetics within and without the context of chemotherapy may give the predictions of the best treatment scenario for maximal suppression of total cancer cells population growth, based on the calculated *underlying field* function behavior. An essential part of this scenario will depend on identifying the treatment time points at which elimination of specific underlying field factors will be predicted as crucial for cancer population abolishment.

## References

1. Reya T., Morrison S.J., Clarke M.F., Weissman I.L. Stem cells, cancer, and cancer stem cells. Nature, 414 (2001), 105–111.

2. Dean M., Fojo T. and Bates S. Tumour stem cells and drug resistance. Nat. Rev. Cancer, 5(2005), 275–284.

3. Clarke M.F., Dick J.E., Dirks P.B., Eaves C.J., Jamieson C.H., Jones D.L., Visvader J., Weissman I.L., Wahl, G.M. Cancer stem cells-Perspectives on current status and future directions: AACR workshop on cancer stem cells. Cancer Res., 66(2006), 9339–9344.

4. Bao S., Wu Q., McLendon R.E., Hao Y., Shi Q., Hjelmeland A.B., Dewhirst M.W., Bigner D.D., Rich J.N. Glioma stem cells promote radioresistance by preferential ac-tivation of the DNA damage response. Nature, 444 (2006), 756–760.

5. O’Brien C.A., Pollett A., Gallinger S., Dick J.E. A human colon cancer cell capable of initiating tumour growth in immunodeficient mice. Nature, 445 (2007), 106–10.

6. Ricci-Vitiani L., Lombardi D.G., Pilozzi E., Biffoni M., Todaro M., Peschle C., DeMaria, R. Identification and expansion of human colon-cancer-initiating cells. Nature, 445 (2007), 111–115.

7. Li C., Heidt D.G., Dalerba P., Burant C.F., Zhang L., Adsay V., Wicha M., Clarke M.F., Simeone D.M. Identification of pancreatic cancer stem cells. Cancer Res., 67 (2007), 1030–7.

8. Mani S.A., Guo W., Liao M.J., Eaton E.N., Ayyanan A., Zhou A.Y. et al. The epithelial-mesenchymal transition generates cells with properties of stem cells. Cell, 133 (2008), 704–715.

9. Ginestier C., Wicha M.S. Mammary stem cell number as a determinate of breast cancer risk. Breast Cancer Res., 9 (2007), 109–126.

10. Cicalese A., Bonizzi G., Pasi C.E., Faretta M., S. Ronzoni et al. The Tumor Suppressor p53 Regulates Polarity of Self-Renewing Divisions in Mammary Stem Cells. Cell, V. 138, Iss.6, 2009, 1083–1095.

11. Zhang S., Balch C., Chan M.W., Lai H.C., Matei D., Schilder J.M. et al. Identification and characterization of ovarian cancer-initiating cells from human tumors. Cancer Res., 68 (2008), 4311–20.

12. Maitland N.J. Collins A.T. Prostate cancer stem cells: a new target for therapy. J. Clin.Oncol., 26 (2008), 2862–70.

13. Gupta P.B., Onder T.T., Jiang G., Tao K., Kuperwasser C., Weinberg R.A. Lander E.S. Identification of selective inhibitors of cancer stem cells by high-throughput screening. Cell, 138(2009), 645–59.

14. Diehn M., Clarke M.F. Cancer stem cells and radiotherapy: new insights into tumor radioresistance. J. Natl. Cancer Inst., 98 (2006), 1755–1757.

15. Loeffler M., Roeder I. Conceptual models to understand tissue stem cell organization. Current Opinion in Hematology 2004, 11:81–87.

16. Roeder I., Herberg M., Horn M. An “Age” structured model of hemapoietic stem cell organization with application to chronic myeloid leukemia. Bull. Math. Biol., 71 (2009), 602–626.

17. D’Onofrio A., Tomlison I.P.M. A nonlinear mathematical model of cell renewal, turnover and tumorigenesys in colon crypts. J. Theor. Biol., 244 (2007), 367–374.

18. Chaffer C.L., Brueckmann I., Scheel C., Kaestli A.J., Wiggins P.A., Rodrigues L.O., et al. Normal and neoplastic nonstem cells can spontaneously convert to a stem-like state. Proc. Natl. Acad. Sci. 108, (2011) 7950–7955.

19. Gupta PB, Fillmore CM, Jiang G, Shapira SD, Tao K, Kuperwasser C, Lander ES. Stochastic state transitions give rise to phenotypic equilibrium in populations of cancer cells. Cell. (2011) Aug 19; 146(4):633–44.

20. Enderling H., Hlatky L., Hahnfeldt P. Cancer Stem Cells: A Minor Cancer Subpopulation that Redefines Global Cancer Features. Front Oncol. 2013; 3: 76.

21. Goldman et al., Nature communications, 6:6139, 2014.

22. Johnston M.D., Edwards C.M., Bodmer W.F., Maini P.K., Chapman, S.J. Mathematical modelling of cell population dynamics in the colonic crypt and in colorectal cancer. PNAS, 104 (2007), 4008–4013.

23. Theise N.D., d'Inverno M. Understanding cell lineages as complex adaptive systems. Blood Cells Mol Dis. 2004 Jan-Feb;32(1):17–20.

24. Theise N.D. Perspective: stem cells react! Cell lineages as complex adaptivesystems. Exp Hematol 2004, 32:25–27.

25. Michor F., Mathematical models of cancer stem cells. J. Clin. Oncol., 26 (2008), 2854–2861.

26. Beretta E, Capasso V, Morozova N. Mathematical Modelling of Cancer Stem Cells Population Behavior. Math Mod Nat Phen, V. 7, Is. 1, p.279–305 (2012)

27. Beretta E, Capasso V, Harel-Bellan A, Morozova N. Some Results on the Population Behavior of Cancer Stem Cells. In: New Challenges for Cancer Systems Biomedicine. d’Onofrio, Cerrai, Gandolfi (eds), Simai Springer Series. 2012

